# Assessing Autophagy Flux in Glioblastoma Temozolomide Resistant Cells

**DOI:** 10.1101/2024.08.09.607348

**Authors:** Courtney Clark, Amir Barzegar-Behrooz, Marco Cordani, Shahla Shojaei, Saeid Ghavami

## Abstract

Autophagy is a critical cellular process involved in the degradation and recycling of cytoplasmic components, playing a dual role in cancer by either promoting cell survival or facilitating cell death. In glioblastoma (GB), autophagy has been implicated in resistance to the chemotherapeutic agent Temozolomide (TMZ). This study presents a novel method to accurately measure autophagy flux in TMZ-resistant glioblastoma cells, combining advanced imaging techniques with biochemical assays. By quantifying key autophagy markers such as LC3-II and SQSTM1, our approach provides detailed insights into the dynamic processes of autophagosome formation and clearance under therapeutic stress. This method not only advances our understanding of autophagy in GB chemoresistance but also has significant implications for the development of autophagy-targeted therapies. The ability to monitor and manipulate autophagy flux in real-time offers a promising avenue for monitoring and understnading TMZ resistance and improving patient outcomes in glioblastoma treatment.

## 1. Introduction

Autophagy is a fundamental cellular process responsible for the degradation and recycling of cytoplasmic components, including damaged organelles and misfolded proteins, through lysosomal degradation (1-3). This dynamic process, known as autophagy flux, involves the formation of autophagosomes that sequester cellular debris and their subsequent fusion with lysosomes for degradation (4-6). Measuring autophagy flux provides insights into the dynamic nature of autophagy, distinguishing between increased autophagosome formation and impaired autophagosome clearance (7-9). Accurate assessment of autophagy flux is crucial for understanding its role in various physiological and pathological conditions, including cancer (10, 11).

Glioblastoma (GB) is the most aggressive and common primary brain tumor in adults, characterized by rapid growth, extensive infiltration, and resistance to conventional therapies (4, 5, 12). Despite advances in surgical resection, radiotherapy, and chemotherapy, the prognosis for GB patients remains dismal, with a median survival of 12-15 months (13, 14). One of the standard chemotherapeutic agents for GB is Temozolomide (TMZ), an oral alkylating agent that induces DNA damage, leading to cell death. However, the development of TMZ resistance is a significant challenge, limiting its therapeutic efficacy and contributing to GB’s poor prognosis (2, 15, 16).

Emerging evidence suggests that autophagy plays a dual role in cancer, acting as a tumor suppressor by degrading damaged organelles and proteins and as a survival mechanism by providing nutrients through the recycling of cellular components under stress conditions (17-19). In GB, autophagy has been implicated in promoting resistance to TMZ, allowing cancer cells to survive and adapt to chemotherapy-induced stress (9, 20, 21). Autophagy flux in GB TMZ-resistant cells is a dynamic process that contributes to the maintenance of cellular homeostasis and the survival of cancer cells under therapeutic pressure (22, 23).

Several studies have highlighted the role of autophagy flux in GB TMZ resistance, demonstrating that autophagy inhibitors can sensitize resistant GB cells to TMZ, thereby enhancing its cytotoxic effects (4, 5). This indicates that autophagy flux is a critical adaptive response that enables GB cells to withstand TMZ-induced stress (10, 24). Furthermore, autophagy-related genes (ATGs) and proteins involved in the autophagy pathway, such as Beclin-1, LC3, and SQSTM1, are often dysregulated in GB, contributing to the modulation of autophagy flux and chemoresistance (17). Understanding the mechanisms underlying autophagy flux in GB TMZ-resistant cells is essential for developing novel therapeutic strategies to overcome resistance and improve patient outcomes (23).

Our novel method for measuring autophagy flux in GB TMZ-resistant cells provides a comprehensive approach to quantifying the dynamic process of autophagy. This method involves advanced imaging techniques, such as confocal and transmission electron microscopy, combined with biochemical assays to monitor the levels of autophagy markers, including LC3-II and SQSTM1, under various treatment conditions. By integrating these techniques, we can accurately assess the formation and clearance of autophagosomes, providing a detailed understanding of autophagy flux in response to TMZ treatment. This method applies to basic science research and has significant implications for clinical scientists seeking to develop autophagy-targeted therapies for GB.

In summary, the ability to measure autophagy flux in GB TMZ-resistant cells is crucial for elucidating the role of autophagy in chemoresistance and identifying potential therapeutic targets. Our method offers a robust and reliable approach to quantify autophagy flux, contributing to the advancement of both basic and clinical research in GB. By providing detailed insights into the autophagic process, this method can facilitate the development of novel strategies to overcome TMZ resistance, ultimately improving the therapeutic outcomes for GB patients.

## 2. Detailed Method for Measuring Autophagy Flux in U251 TMZR and U251 TMZ NR Cells Using Immunoblotting

### 2.1 Materials and Reagents

1. U251 TMZ-resistant (TMZR) and U251 TMZ non-resistant (TMZ NR) cells
2. DMEM high glucose medium (CORNING; Cat#: 50–003-PB)
3. Fetal bovine serum (FBS) (Gibco™; Cat#: 16000044)
4. Penicillin-streptomycin (Gibco™; Cat#: 41400045)
5. Temozolomide (TMZ) (CAS Number: 85622-93-1)
6. Phosphate Buffered Saline (PBS) (Sigma-Aldrich, Cat #D8537)
7. Trypsin-EDTA (GIBCO® (Thermo Fisher Scientific, MA, USA)
8. Lysis buffer: 20 mM Tris-HCl (pH 7.5), 0.5 mM PMSF, 0.5% Nonidet P-40, 100 μM β-glycerol 3-phosphate, 0.5% protease and phosphatase inhibitor cocktail
9. Lowry protein assay kit (Bio-Rad, Oakville, ON, CA)
10. SDS-PAGE reagents (Bio-Rad, Oakville, ON, CA): Acrylamide/Bis-acrylamide Solution: (For gel polymerization), SDS (Sodium Dodecyl Sulfate): (Denaturing agent), Tris Buffer: (Maintains pH), APS (Ammonium Persulfate): (Polymerization initiator), TEMED (Tetramethylethylenediamine): (Polymerization accelerator), Running Buffer: (Contains Tris, glycine, and SDS), Loading Buffer: Contains SDS, bromophenol blue, glycerol, and β-mercaptoethanol or DTT, Protein Ladder/Marker: For molecular weight reference, Sample Buffer: Typically contains SDS, Tris, glycerol, and a reducing agent.
11. Polyvinylidene difluoride (PVDF) membranes (Thermo Fisher Scientific, Waltham, MA, USA; Cat#: 88518
12. Blocking solution: 5% skim milk in TBST (1X tris-buffered saline, 0.025% Tween 20) (Sigma, St. Louis, MO, USA)
13. Primary antibodies: LC3 (Sigma Aldrich, Oakville, CA; Cat#: L7543, 1:2500 dilution), SQSTM1 (Cell Signaling Technology, Canada; Cat#: 5114, 1:1000 dilution)
14. Secondary antibodies: Fluorochrome-conjugated secondary antibodies (Alexa Fluor 488 (Thermo Fisher Scientific, Waltham, A, USA; Cat#: A21206) and Alexa Fluor 594 (Thermo Fisher Scientific, Waltham, MA, USA; Cat#: R37119).
15. Enhanced chemiluminescence (ECL) detection kit (Amersham-Pharmacia Biotech)
16. Bafilomycin A1 (Sigma Aldrich, Cat#B1793-10UG)

### 2.2. Methods

#### 2.2.1. Cell Culture

##### Culturing U251 Cells

1. Prepare DMEM high glucose medium supplemented with 10% FBS and 1% penicillin-streptomycin. **See Notes and Tips 1**.
2. Thaw U251 TMZR and U251 TMZ NR cells and transfer them to T75 culture flasks containing 20 mL of prepared DMEM medium.
3. Maintain U251 TMZR cells in DMEM medium with 250 μM TMZ. Incubate all flasks at 37°C in a 5% CO2 humidified incubator.
4. Change the medium every 2-3 days. For TMZR cells, remove TMZ-containing medium and replace it with TMZ-free DMEM medium 24 hours before collection to mimic withdrawal conditions. **See Notes and Tips 2**.

### 2.3. Cell Passaging and Collection

#### 2.3.1. Cell Passaging

1. When cells reach 80-90% confluency, aspirate the medium and wash cells with 5 mL PBS. **See Notes and Tips 3**.
2. Add 3 mL of trypsin-EDTA solution to the flask and incubate at 37°C for 2-3 minutes until cells detach. **See Notes and Tips 4**.
3. Neutralize trypsin by adding 7 mL of DMEM medium containing FBS.
4. Transfer cell suspension to a 15 mL centrifuge tube and centrifuge at 300 x g for 5 minutes. **See Notes and Tips 5**.
5. Aspirate the supernatant and resuspend the cell pellet in a fresh DMEM medium.
6. Seed cells at the appropriate density (e.g., 2 × 10^5 cells/mL) in new T75 flasks or culture dishes for subsequent experiments.

#### 2.3.2. Cell Collection

1. At the designated time point, aspirate the medium and wash cells with 5 mL PBS.
2. Add 3 mL of trypsin-EDTA solution to the flask and incubate at 37°C for 2-3 minutes until cells detach. **See Notes and Tips 4**.
3. Neutralize trypsin by adding 7 mL of DMEM medium containing FBS.
4. Transfer cell suspension to a 15 mL centrifuge tube and centrifuge at 300 x g for 5 minutes. **See Notes and Tips 5**.
5. Wash the cell pellet with 5 mL cold PBS and centrifuge again at 300 x g for 5 minutes. **See Notes and Tips 6 and 7**.
6. Aspirate the supernatant and resuspend the cell pellet in 1 mL cold PBS. **See Notes and Tips 7**.

### 2.4. Preparation of Cell Lysates

#### 2.4.1 Cell Lysis

1. Transfer the resuspended cell pellet to a microcentrifuge tube.
2. Add 200 μL of lysis buffer (20 mM Tris-HCl, pH 7.5, 0.5 mM PMSF, 0.5% Nonidet P-40, 100 μM β-glycerol 3-phosphate, 0.5% protease and phosphatase inhibitor cocktail) to the tube.
3. Sonicate the lysate briefly using a sonicator set to 3 cycles of 5 seconds each, with 10-second intervals on ice, to ensure complete cell disruption. **See Notes and Tips 8**.
4. Centrifuge the lysate at 10,000 x g for 10 minutes at 4°C to remove cell debris.
5. Transfer the clear supernatant (protein lysate) to a new microcentrifuge tube.

#### 2.4.2. Protein Quantification

1. Use the Lowry protein assay kit to determine the protein concentration in the lysate. **See Notes and Tips 9**.
2. Prepare a standard curve using BSA standards ranging from 0 to 1 mg/mL.
3. Add 5 μL of each protein sample or standard to a 96-well plate in triplicate.
4. Add 25 μL of Lowry reagent to each well, mix gently, and incubate for 10 minutes at room temperature.
5. Add 125 μL of Folin-Ciocalteu reagent, mix gently, and incubate for 30 minutes at room temperature.
6. Measure the absorbance at 750 nm using a microplate reader.
7. Calculate the protein concentration of each sample based on the standard curve and adjust the lysate concentration to 1 mg/mL using lysis buffer.

### 2.5. Immunoblotting

#### 2.5.1. SDS-PAGE and Transfer

1. Prepare a 10-15% polyacrylamide gel for SDS-PAGE.
2. Load 30 μg of protein from each sample into the wells and run the gel at 100 V for 1.5 hours.
3. Transfer the separated proteins to PVDF membranes at 100 V for 2 hours under reducing conditions.

#### 2.5.2. Blocking and Primary Antibody Incubation

1. Block the membranes with 5% skim milk in TBST for 1 hour at room temperature. **See Notes and Tips 10**.
2. Incubate membranes overnight at 4°C with primary antibodies against LC3 (1:2500 dilution, Sigma) and SQSTM1 (1:1000 dilution, Cell Signaling) in 1% skim milk in TBST.

#### 2.5.3. Secondary Antibody Incubation and Detection

1. Wash membranes with TBST (3 washes, 10 minutes each).
2. Incubate with HRP-conjugated secondary antibodies for 1 hour at room temperature.
3. Wash membranes again with TBST (3 washes, 10 minutes each).
4. Develop the blots using an ECL detection kit and visualize bands using a BioRad imager.

### 2.6. Evaluation of Autophagy Flux

#### 2.6.1. LC3 Lipidation and SQSTM1 Degradation

1. Assess LC3-II levels (lipidated form) and SQSTM1 degradation using immunoblotting. Increased LC3-II indicates autophagosome formation, while decreased SQSTM1 indicates autophagosome degradation.

#### 2.6.2. Bafilomycin A1 Treatment for Flux Assessment

##### Treatment

1. Treat cells with 100 nM Bafilomycin A1 for 6 hours to inhibit autophagosome-lysosome fusion.
2. Include a control group without Bafilomycin A1 treatment for comparison.

##### Harvesting Cells Post-Bafilomycin Treatment

1. Wash cells with PBS and harvest by trypsinization. **See Notes and Tips 6**.
2. Pellet cells by centrifugation at 300 x g for 5 minutes and wash with cold PBS. **See Notes and Tips 7**.

##### Cell Lysis

1. Resuspend the cell pellet in lysis buffer as previously described.
2. Sonicate the lysate briefly (3 cycles of 5 seconds each).
3. Centrifuge the lysate at 10,000 x g for 10 minutes at 4°C.

##### Protein Quantification

1. Use the Lowry protein assay kit to determine protein concentration and adjust to 1 mg/mL. **See Notes and Tips 9**.

##### SDS-PAGE and Transfer

1. Separate proteins on 10-15% SDS-PAGE and transfer to PVDF membranes.

##### Blocking and Antibody Incubation

1. Block with 5% skim milk in TBST.
2. Incubate overnight with primary antibodies (LC3 and SQSTM1) at specified dilutions. **See Notes and Tips 10**.

##### Secondary Antibody and Detection

1. Wash membranes with TBST, incubate with HRP-conjugated secondary antibodies.
2. Develop blots using ECL detection and visualize bands with a BioRad imager.

### 2.7. Detailed Evaluation of Autophagy Flux

#### 2.7.1. Baseline Autophagy Measurement

1. Measure LC3-II and SQSTM1 levels in U251 TMZR and TMZ NR cells under normal conditions.

#### 2.7.2. Autophagy Inhibition with Bafilomycin A1

1. Treat cells with 100 nM Bafilomycin A1 for 6 hours.
2. Harvest cells, lyse, and prepare protein samples as described.
3. Perform immunoblotting to detect LC3-II and SQSTM1 levels.

#### 2.7.3. Comparison and Interpretation

1. Compare the levels of LC3-II and SQSTM1 between treated and untreated cells.
2. A significant increase in LC3-II and SQSTM1 levels in Bafilomycin A1 treated cells compared to untreated controls

## 3. Detailed Method for Transmission Electron Microscopy (TEM) and Evaluation of Autophagy Flux

### 3.1. Materials and Reagents

- U251 TMZ-resistant (TMZR) and U251 TMZ non-resistant (TMZ NR) cells
- DMEM high glucose medium (CORNING; Cat#: 50–003-PB)
- Fetal bovine serum (FBS) (Gibco™; Cat#: 16000044)
- Penicillin-streptomycin (Gibco™; Cat#: 41400045)
- Karnovsky fixative
- 5% sucrose in 0.1M Sorenson’s phosphate buffer
- 1% osmium tetroxide in 0.1M Sorenson’s buffer
- Ethanol series for dehydration (50%, 70%, 90%, 100%) (UN 1170, Greenfield Global, # PO25EAAN).
- Embed 812 resin
- Uranyl acetate
- Lead nitrate
- Copper grids (EMS, #G200-Cu)
- Philips CM10 electron microscope

### 3.2. Method

### 3.3. Cell Culture

#### 3.3.1. Culturing U251 Cells

∘ Culture U251 TMZR and U251 TMZ NR cells in DMEM high glucose medium supplemented with 10% FBS and 1% penicillin-streptomycin. **See Notes and Tips 1**.
∘ Incubate cells at 37°C in a 5% CO2 humidified incubator until they reach 80-90% confluency. **See Notes and Tips 3**.

### 3.4. Fixation

#### 3.4.1. Cell Fixation

- Wash cells with PBS and detach using trypsin-EDTA. **See Notes and Tips 6**.
- Pellet cells by centrifugation at 300 x g for 5 minutes. **See Notes and Tips 5**.
- Resuspend the cell pellet in 2 mL of Karnovsky fixative and incubate for 1 hour at 4°C.

#### 3.4.2. Post-Fixation

∘ After fixation, wash cells with 0.1M Sorenson’s phosphate buffer.
∘ Resuspend the pellet in 5% sucrose in 0.1M Sorenson’s phosphate buffer and store overnight at 4°C.

#### 3.4.3. Osmium Tetroxide Post-Fixation

∘ Pellet cells again and fix with 1% osmium tetroxide in 0.1M Sorenson’s buffer for 2 hours at room temperature.

### 3.5. Dehydration and Embedding

#### 3.5.1. Dehydration

- Dehydrate cells through a graded ethanol series: 50%, 70%, 90%, and 100% ethanol, each for 10 minutes.
- Perform final dehydration with 100% ethanol for 20 minutes.

#### 3.5.2. Embedding

∘ Infiltrate cells with Embed 812 resin: first, 1:1 mixture of resin and ethanol for 1 hour, then pure resin overnight at room temperature.
∘ Embed the cells in fresh resin and polymerize at 60°C for 48 hours.

### 3.6. Sectioning and Staining

#### 3.6.1. Sectioning

- Cut thin sections (70-90 nm) using an ultramicrotome.
- Place sections on copper grids.

#### 3.6.2. Staining

∘ Stain sections with 2% uranyl acetate for 10 minutes.
∘ Wash with distilled water and stain with lead citrate for 5 minutes.

#### 3.7. Imaging

#### 3.7.1. Imaging

- Examine the stained sections using a Philips CM10 electron microscope.
- Capture images of representative fields at appropriate magnifications (e.g., 10,000x to 20,000x).

### 3.8. Evaluation of Autophagy Flux

#### 3.8.1. Counting Autophagosomes and Autolysosomes

∘ **Autophagosomes:** Identify and count double-membrane vesicles containing cytoplasmic material (autophagosomes) in the cytosol.
∘ **Autolysosomes:** Identify and count single-membrane vesicles containing partially degraded material (autolysosomes).

#### 3.8.2. Comparison Between Conditions

∘ Compare the number of autophagosomes and autolysosomes in U251 TMZR and TMZ NR cells.
∘ Analyze cells with and without treatment (e.g., Bafilomycin A1) to determine the impact on autophagy flux.

#### 3.8.3. Data Interpretation

∘ Increased number of autophagosomes in the presence of Bafilomycin A1 indicates blocked autophagosome-lysosome fusion, reflecting active autophagosome formation.
∘ A decrease in autophagosome numbers with a corresponding increase in autolysosomes indicates effective autophagosome-lysosome fusion and subsequent degradation.

#### 3.8.4. Quantitative Analysis

∘ Calculate the average number of autophagosomes and autolysosomes per cell.
∘ Use statistical analysis to compare the differences between treated and untreated cells, and between TMZR and TMZ NR cell lines.

This detailed TEM method provides a comprehensive approach to evaluate autophagy flux by analyzing the numbers of autophagosomes and autolysosomes, offering insights into the autophagic activity in glioblastoma cells (Scheme 1).

## 4. Evaluation of Autophagy Flux in U251 TMZ-NR and TMZ-R Cells

### 4.1. Expression of LC3-beta II (LC3 lipidation) and SQSTM1 degradation in Immunoblotting

Autophagy flux was assessed by measuring the expression levels of LC3-beta II (LC3 lipidation) and SQSTM1 degradation using immunoblotting. The microtubule-associated proteins 1A/1B light chain 3B (LC3) is essential in autophagy, undergoing lipidation to form LC3-II, which is associated with autophagosome membranes. SQSTM1 is a selective substrate of autophagy, degraded within autolysosomes, serving as a marker for autophagic degradation.

#### 4.1.2. Step I: Baseline Autophagy Flux

In the baseline scenario, without any treatments, we compared the levels of LC3-beta II and SQSTM1 in U251 TMZ-NR and TMZ-R cells:

- **U251 TMZ-NR Cells**: Higher levels of LC3-beta II and lower levels of SQSTM1 indicate active autophagy flux.
- **U251 TMZ-R Cells**: Elevated levels of both LC3-beta II and SQSTM1 suggest impaired autophagy flux, potentially due to defective autophagosome-lysosome fusion or lysosomal function.

### 4.2. Number of Cytosolic Autophagosomes and Autolysosomes in Transmission Electron Microscopy (TEM)

Transmission electron microscopy (TEM) was employed to visually quantify the number of cytosolic autophagosomes and autolysosomes in U251 TMZ-NR and TMZ-R cells. This provided structural evidence supporting the biochemical data from immunoblotting.

#### 4.2.1. Step II: TEM Analysis

- **U251 TMZ-NR Cells**: TEM images revealed a higher number of autolysosomes compared to autophagosomes, consistent with active autophagy flux where autophagosomes are rapidly fused with lysosomes for degradation.
- **U251 TMZ-R Cells**: TEM images showed an accumulation of autophagosomes with fewer autolysosomes, indicating a block in the autophagic process, likely at the autophagosome-lysosome fusion stage.

### 4.3. Evaluation of Autophagy Flux using Bafilomycin A1

To further confirm the autophagy flux status, we used Bafilomycin A1 (BafA1), an inhibitor of vacuolar-type H+-ATPase, which prevents the acidification of lysosomes and inhibits autophagosome-lysosome fusion.

#### 4.3.1. Step III: Immunoblotting with Bafilomycin A1

We treated U251 TMZ-NR and TMZ-R cells with Bafilomycin A1 and evaluated the changes in LC3-beta II and SQSTM1 levels.

- **U251 TMZ-NR Cells**: Upon BafA1 treatment, there was a significant accumulation of LC3-beta II and SQSTM1, indicating that the autophagic flux is active under normal conditions and BafA1 effectively blocks the final degradation step.
- **U251 TMZ-R Cells**: BafA1 treatment did not result in a significant increase in LC3-beta II and SQSTM1 levels, suggesting that autophagy flux is already impaired at an earlier stage, before the autophagosome-lysosome fusion.

**The overall evaluation of autophogy flux in U251 TMZ NR and R cells has been summarized in Scheme 2 and was experimentally showed in Figure-1**.

**Figure 1.**
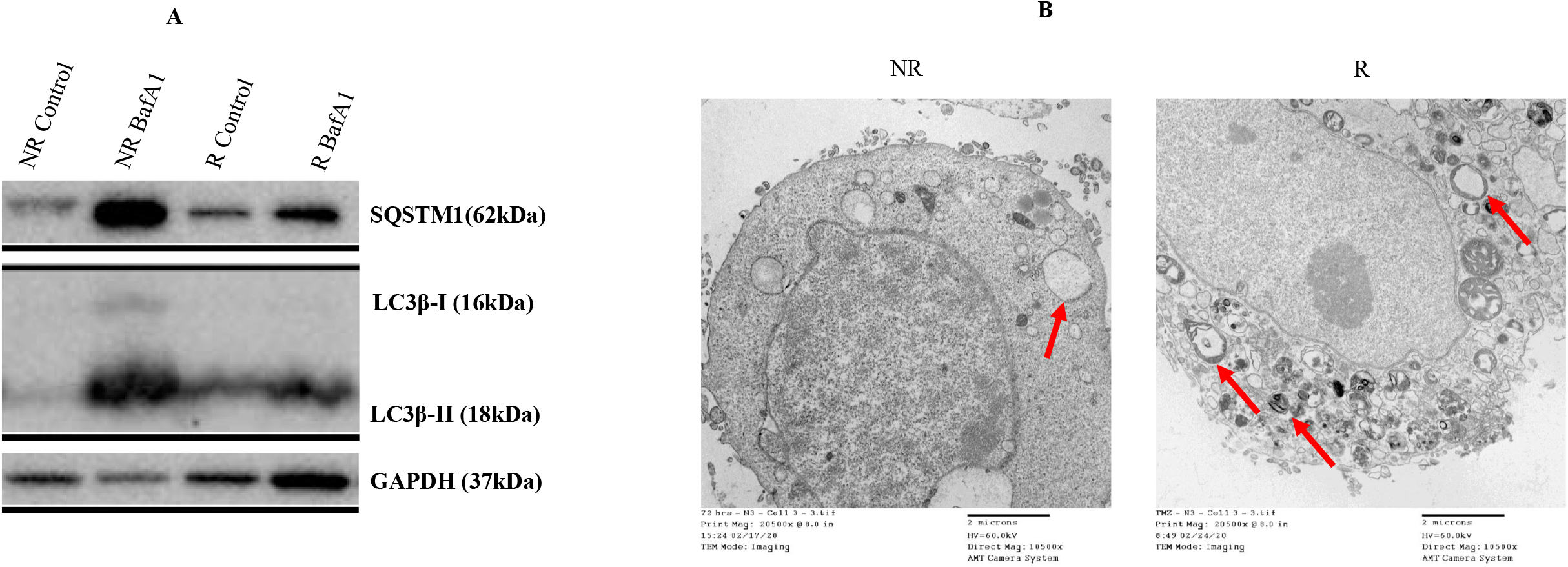
Autophagy flux is inhibited in U251 TMZ-Chemoresistant GB cells.. U251 TMZ-NR and R GB cells were treated with Bafilomycin-A1 (5nM) for 72 hours. Autophagy was assessed via immunoblotting, evaluating SQSTM1 degradation, LC3β lipidation, and LC3βII formation, with GAPDH as a loading control. Bafilomycin-A1 inhibited autophagy flux in U251 TMZ-NR cells but had no significant effect on U251 TMZ-R cells. **B**. U251 TMZ-NR and R GB cells were cultured for 72 hours and prepared for TEM (20,500x magnification) to assess autophagy by counting cytosolic double-membrane vacuoles. The results showed a higher number of double-membrane vacuoles in U251 TMZ-R cells compared to U251 TMZ-NR cells.

## 5. Notes and Tips

Here are ten important notes and tips for achieving optimal results.

1. **Consistent Medium Preparation:** Ensure that the DMEM high glucose medium is consistently supplemented with 10% FBS and 1% penicillin-streptomycin. This is crucial for maintaining the health and growth of U251 cells.
2. **TMZ Treatment Specificity:** For U251 TMZR cells, always maintain them in DMEM medium with 250 μM TMZ. Remember to remove TMZ-containing medium and replace it with TMZ-free DMEM medium 24 hours before collection to mimic withdrawal conditions accurately.
3. **Cell Confluency Check:** Regularly monitor cell confluency, ensuring cells are passaged at 80-90% confluency. Over-confluent or under-confluent cultures can affect cell behavior and experimental results.
4. **Trypsinization Timing:** Be precise with the trypsin-EDTA incubation time (2-3 minutes). Over-trypsinization can damage cells, while under-trypsinization may not detach cells effectively.
5. **Centrifugation Consistency:** Maintain consistent centrifugation conditions (300 x g for 5 minutes). Variations can result in incomplete cell pelleting or cell damage.
6. **PBS Washing:** Thoroughly wash cells with PBS to remove any residual medium or trypsin before proceeding to the next steps, such as passaging or collection.
7. **Cold PBS for Cell Collection:** Use cold PBS for washing and resuspending cell pellets during cell collection to minimize cell metabolism and preserve cellular integrity for downstream applications.
8. **Sonication for Complete Disruption:** When sonicating the lysate, ensure the sonicator is set to 3 cycles of 5 seconds each with 10-second intervals on ice. This helps in complete cell disruption without overheating the samples.
9. **Accurate Protein Quantification:** Use the Lowry protein assay kit with a carefully prepared standard curve to accurately determine protein concentrations. Inaccurate quantification can lead to errors in downstream applications like immunoblotting.
10. **Controlled Blocking and Antibody Incubation:** During immunoblotting, ensure blocking of membranes is done with 5% skim milk in TBST for exactly 1 hour, and primary antibody incubation is done overnight at 4°C. These steps are critical for reducing background noise and ensuring specific binding of antibodies.

## Acknowledgement

SG conceptualized and designed the study. CC performed the experiments. SG, SS, MC, SS, ABB analyzed the data. SG and MC wrote the manuscript. SG, CC, SS, ABB and MC reviewed and edited the manuscript. SS developed GB TMR R cells. SG funded supervised the experiments. The authors acknowledge Simone C da Silva Rosa for the ICC evaluation of autophagy.

## Scheme Legends

**Scheme 1:**
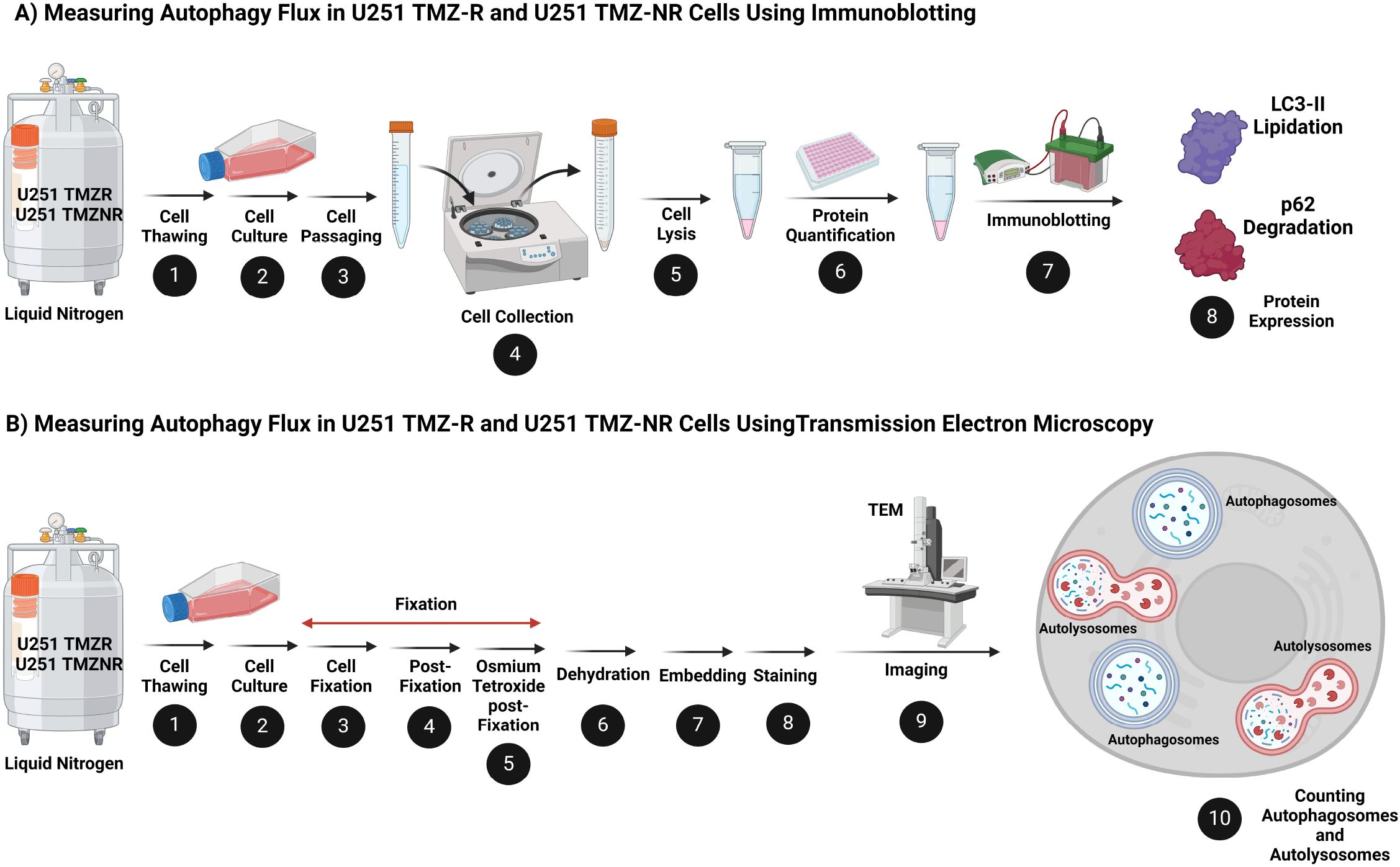
Overview of the methodology used to measure autophagy flux in U251 TMZ-resistant (TMZR) and non-resistant (TMZ NR) cells. (A) Immunoblotting: The levels of autophagy markers LC3-II and SQSTM1 are quantified to assess autophagosome formation and degradation. (B) Electron microscopy: Ultrastructural analysis is performed to visualize and count autophagosomes and autolysosomes, providing detailed insights into the autophagic process at a cellular level. This combined approach offers a comprehensive evaluation of autophagy flux, crucial for understanding the mechanisms of TMZ resistance in glioblastoma cells.

**Scheme 2:**
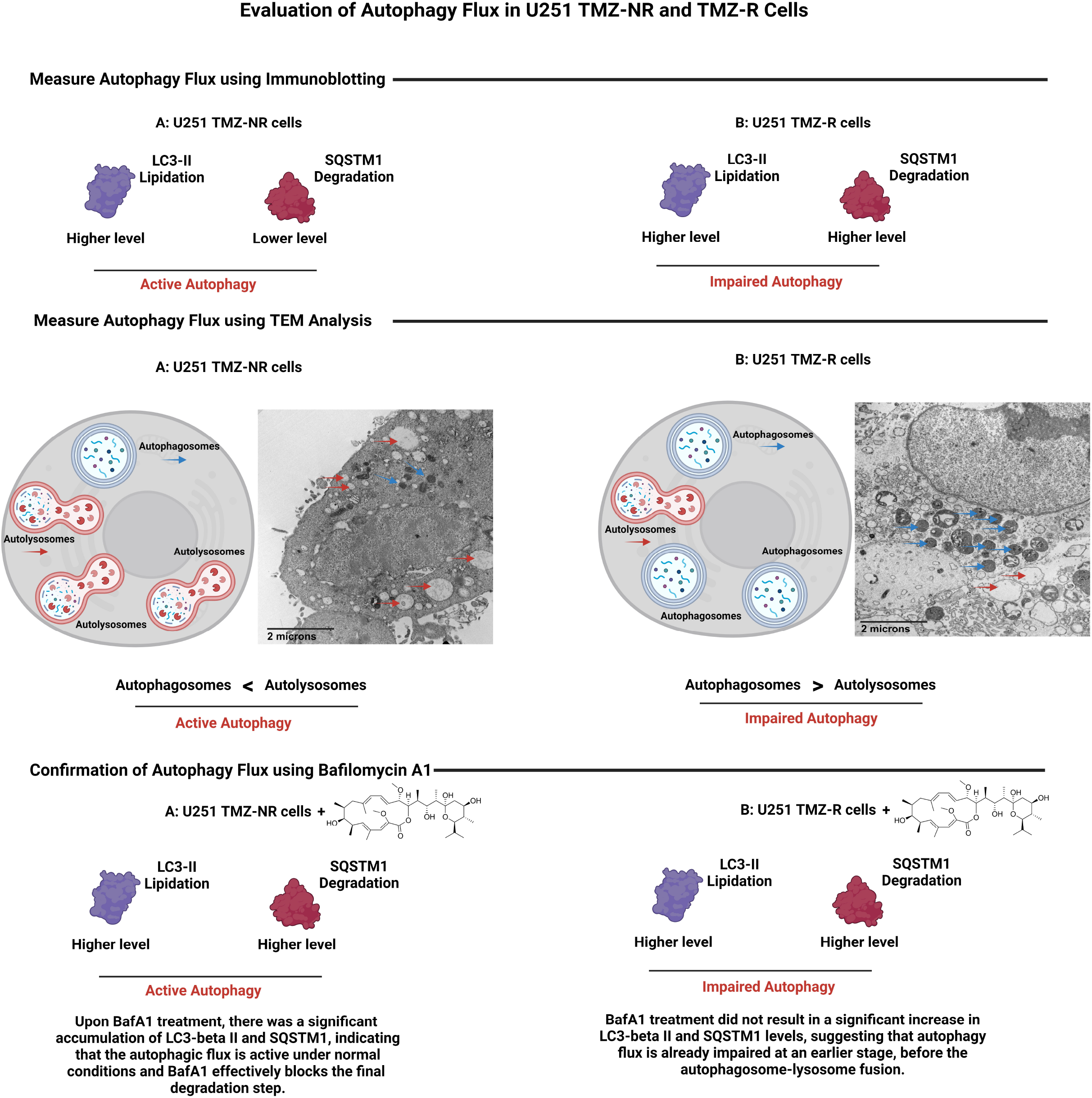
Steps in the confirmation of autophagy flux. (A) Immunoblotting: The levels of LC3-II and SQSTM1 are measured to evaluate autophagosome degradation and autophagy flux. (B) Transmission Electron Microscopy (TEM): The numbers of autophagosomes and autolysosomes are quantified to assess autophagy flux at the ultrastructural level. (C) Bafilomycin A1 Treatment: Final confirmation of autophagy flux is achieved by evaluating the accumulation of LC3-II and SQSTM1 after treatment, indicating blocked autophagosome-lysosome fusion. This multi-step approach provides a robust evaluation of autophagy flux in U251 cells.

